# Planetary-scale heterotrophic microbial community modeling assesses metabolic synergy and viral impacts

**DOI:** 10.1101/2025.02.13.638167

**Authors:** A. Régimbeau, F. Tian, G. Smith, V J. Riddell, C M. Andreani-Gerard, P. Bordron, M. Budinich, C. Howard-Varona, A. Larhlimi, E. Ser-Giacomi, C. Trottier, L. Guidi, S.J. Hallam, D. Iudicone, E. Karsenti, A. Maass, M.B. Sullivan, D. Eveillard

## Abstract

The oceans buffer against climate change via biogeochemical cycles underpinned by microbial metabolic activities. While planetary-scale surveys provide baseline microbiome data, inferring metabolic and biogeochemical impacts remains challenging. Genome-scale modeling has addressed analogous issues at the cellular level, highlighting key metabolic reactions contingent upon specific ‘environmental’ conditions. Here we adapt this mechanistic modeling framework towards analyzing global ocean microbial communities to reveal metabolic processes predicted to maintain ecosystem functioning. To achieve this, we developed a genome-scale ‘superorganism’ metabolic model for each TARA Ocean metagenome or metatranscriptome (i.e., limited to reactions known from heterotrophic prokaryotes and viruses), and evaluated these models to establish a community-wide ‘metabolic phenotype’ for each sample. To validate, we showed that even with reaction-mappable genes only (∼1/4 of the total genes), model composition revealed metabolism-inferred ecological zones that matched taxonomy-inferred zones. Model inferred metabolic phenotypes revealed reaction cooperation associated with microbial metabolism and organism diversity. These phenotypes also suggest elevated ecological roles for viruses as model predictions suggest they genomically target community-critical metabolic reactions that underpin metabolic phenotype stability, and also demonstrate that, as metabolites are better understood, immediate estimates could be made for where viruses remineralize versus sink carbon. While this new constraints-based, agile, and mechanistic modeling framework is highly upgradable, it already begins to convert molecular-scale environmental omics data to ecological and even planetary-scale biogeochemical features that will better bring microbes and their viruses into Earth system and climate models.

---

Connecting “omic” data (DNA, RNA, protein, and metabolites) to ecosystem-level nutrient fluxes and energy flows is a grand challenge in the life sciences^1–3^. Leveraging petascale microbiome data, a fast-growing microbial metabolic knowledgebase^4,5^, and matched physical and chemical information provides opportunities to develop constrained models to interrogate active metabolic processes. However, this will require new modeling approaches as current options cannot yet simultaneously model myriad taxa across the many orders of magnitude of biological organization (e.g., individual, population, and community levels) required to establish ecosystem-, regional- and even planetary-scale predictions.

Fortunately, advances in several areas offer promise as follows. *First*, the rapid growth in genome-resolved ‘-omics’ observations makes them increasingly available to parameterize large-scale biogeochemical models^6^ and simulate ocean ecosystem behavior^7^. *Second*, community-level gene expression data (metatranscriptomes) can now be used to identify transcribed genes that predict regional ocean response patterns^8^. *Third*, metabolic mapping ^9^ and graph-based^10,11^ approaches – even driven simply by reaction presence or absence – are increasingly used to analyze compositional data^12^ or semi-quantitative read counts for inferring organismal or multi-organismal metabolic phenotypes^13,14^. *Finally*, metabolic modeling has started using gene and reaction lists to tackle ecosystem-level questions^15^ where models were effective for assessing mutualistic interactions^16^, distinguishing competitive or cooperative communities^17,18^, and exploring trade-offs between biological objectives that biological systems aim to maximize (e.g., growth, energy storage or consumption) for each member of the community^19^. Currently, the many scales and approaches in modeling offer tremendous promise for opening new windows into biology, but leave benchmarking and validation requirements highly varied and often incredibly challenging, if not impossible, at scales out of sync with observations^20^.

Here, we sought to build upon these efforts by using metabolic modeling, in particular Genome-Scale Models (GSMs), to mechanistically connect microbial metabolic abilities to biogeochemical processes at ecosystem scales. To build and study these GSMs, we chose the superorganism approach^21,22^, whereby all genes in a given sample are considered part of one “community cell”. This simplification was chosen because any taxonomy-constrained modeling framework suffers from uncertainties about how to model interacting species^23^, the low quantity of prokaryotic reads recovered by Metagenome Assembled Genomes (i.e., from 14 to 19%^24^), and computational processing limitations. From this superorganism choice, metagenomic and metatranscriptomic samples represent the gene content or gene expression, respectively, of a superorganism that was then used to reconstruct metabolic networks via a standard top-down approach^25^ that reduces a “universal” metabolic network by removing reactions that lack genetic evidence (see **Methods** for details). Application to *Tara* Oceans data^8,26^ that spanned multiple depths from 62 stations sampled throughout the global ocean (127 prokaryote-enriched metagenomes paired with 94 prokaryote-enriched metatranscriptomes and informed by virus data from 93 virus-enriched metagenomes; **Fig. 1a**, see **Methods**, and **Supplementary Text 1**) resulted in 221 *Tara* Oceans Genome-Scale Metabolic Models (herein TOGSMMs)..

**Figure 1.**
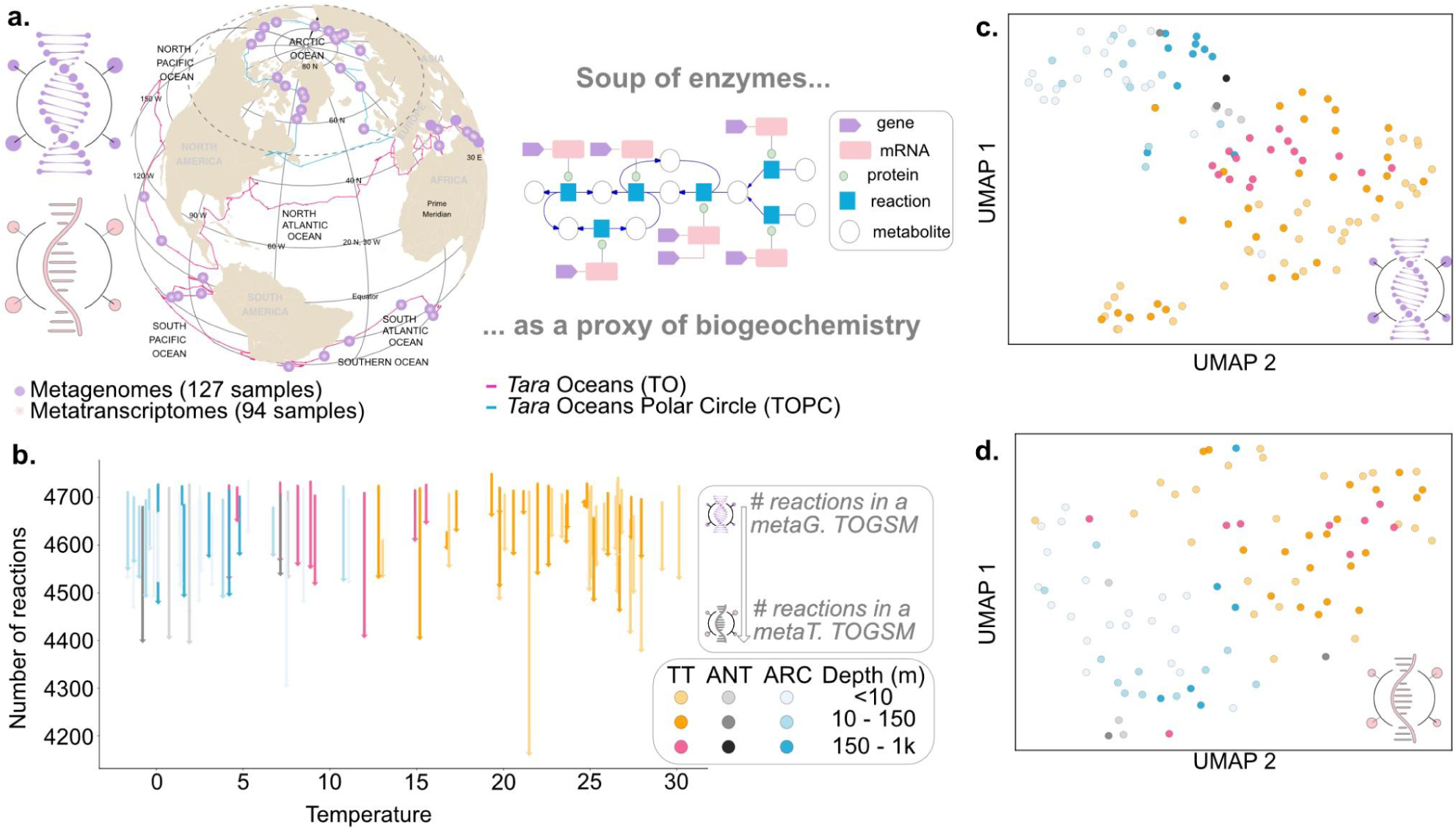
Construction and ecological exploration of TOGSMMs. **a.** Graphical abstract of genome-scale metabolic model reconstruction applied on globally distributed metagenomes (n=127) and metatranscriptomes (n=94) (left). For this, known physics and chemistry of metabolic reactions are captured into metabolic networks (schematic right, where genes are depicted in purple, enzymes in pink, and metabolic reactions in blue that transform metabolites in white). **b.** The number of reactions for each metabolic model was reconstructed per sample from its metagenome, and the matching metatranscriptome was plotted against temperature to assess potential bias due to the latitudinal gradient. In all cases, metagenome-based model are larger (contains more reactions) than their corresponding metatranscriptome-based model, which was interpreted as the per-sample genetic potential (metagenome) versus genetic expression (metatranscriptome) sampled functional repertoires, respectively. Color coding represents previous ecological zones defined by microbial and virus community data^27,26^ (TT = Temperate and Tropical, ANT = Antarctic, ARC = Arctic), and the degree of color represents sampling depth as indicated in the figure. **c** and **d.** Ordination (UMAP, or Uniform Manifold Approximation and Projection) of per-sample presence/absence reaction predictions show that genome-scale metabolic models generated from metagenomes (c) and metatranscriptomes (d) structure along known ecological zones. Colors are the same as **b.**

Even though community metabolic measurements are impossible at scale, we sought to validate this approach by assessing whether TOGSMMs could recover known biological and ecological features. *First*, we observed that per-sample metagenome-constructed TOGSMMs contained more reactions with genetic evidence than metatranscriptome-constructed TOGSMMs, with an average of 4,058 (+/- 28 stdev.) and 3,900 (+/- 79 stdev.) reactions, respectively (**Fig. 1b** and **Supplementary Text 2** for further discussion). These findings are consistent with a community’s expressed transcripts sampled once (i.e., the metatranscriptome) representing only the active subset of genes in the fuller genomic repertoire (i.e., the metagenome). *Second*, despite only a subset of genetic data being incorporated into the model (26% of the total genes observed were mappable to metabolic reactions), TOGSMM-derived ordinations (**Fig. 1c** and **Fig. 1d**) sorted samples into the same global ocean ecological zones known from taxonomy^26,28,29^ or genome content (see **Supplementary Text 2**). Though only a first-level positive sign, this comfortingly showed that the modeled superorganism metabolism indeed follows broader metagenome-inferred ecological patterns. As proposed in the previous *Tara* Ocean analysis^8^, despite not considering autotrophic and eukaryotic communities, the modeled heterotrophic prokaryotic communities reflect a meta-genomic and meta-transcriptomic composition associated with distinct abiotic and biotic environments.

Given the above models’ composition validations, we sought to explore how such modeling could improve our understanding of ocean microbial communities via the identification of crucial metabolic processes necessary for the maintenance of microbial ecosystems. To do this, we sought to provide a mathematical representation for each sample’s observed sequence-related community-wide heterotrophic metabolism, which we term a sample’s ‘metabolic phenotype’. Problematically, solving TOGSMMs would require computing all mathematically feasible reaction fluxes that satisfy the modeling constraints (**Fig. 2a**), which, considering more than four thousand reactions per sample, would result in a highly multidimensional solution space (its dimensions are equal to the number of reactions present in the GSM) that is impossible to solve exhaustively. Instead, we approximated this solution space through metabolic flux sampling analysis^31^ that performs random walks (we conducted 100,000 per sample) through the solution space (**Fig. 2a**, see **Methods** for details).

**Figure 2.**
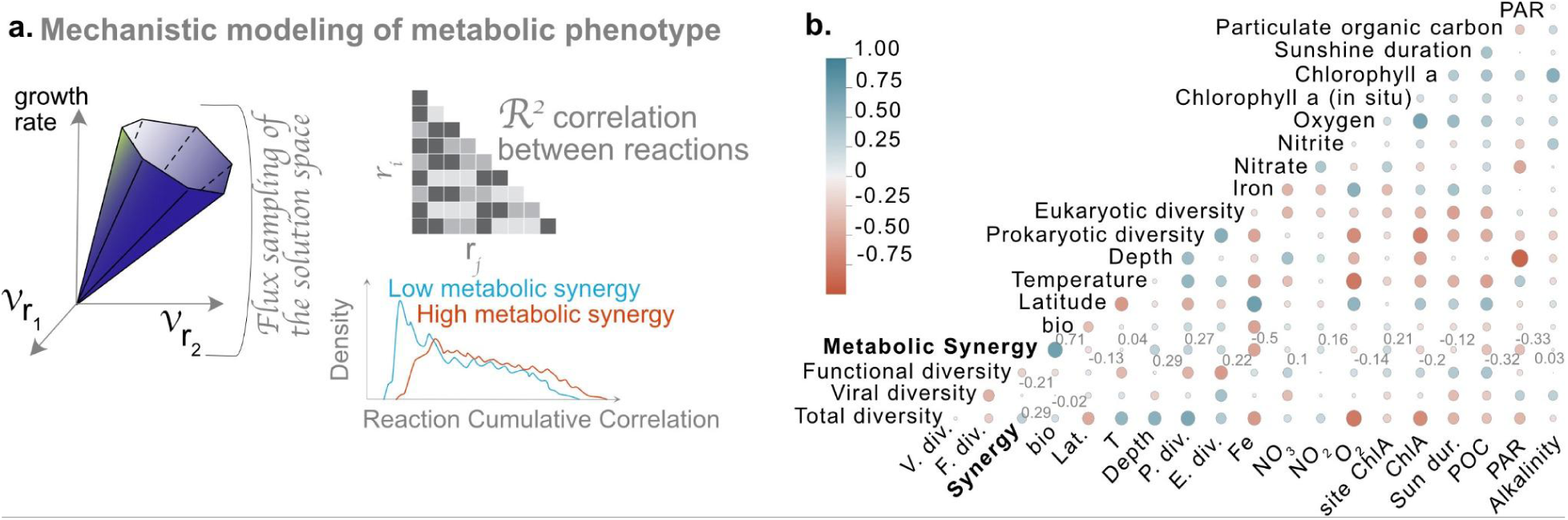
Computing metabolic synergy across the global ocean and comparing other parameters. **a.** Graphical abstract to capture TOGSMM workflows. For each TOGSMM, the metabolic solution space reflects all mechanistic solutions for any environmental conditions (i.e., the fundamental metabolic niche of the community^30^, which we suggest here represents a ‘metabolic phenotype’ for each sample). Because of the complexity of the mechanistic solutions, we summarize them by a flux sampling procedure that estimates each metabolic reaction’s flux distribution. To estimate the reaction’s strength for controlling the metabolic phenotype, pairwise correlations (all reactions vs all other reactions) were computed for each reaction within a network and summarized as a Reaction Cumulative Correlation (RCC) as done elsewhere^30^. For a given network, the RCCs’ distribution determines its metabolic synergy, offering a quantitative score for comparing metabolic phenotypes. **b.** Correlogram of metabolic synergy and environmental variables to assess statistically significant correlations (Spearman correlation).

From these empirical solution space estimates of each TOGSMM, we calculated two metrics. *First*, for each reaction in a TOGSMM, we calculated its Reaction Cumulative Correlation (RCC). This well-known metric^32^ estimates each reaction’s ability to quantitatively and mechanistically impact the overall metabolic reaction network and is calculated as the sum of a reaction’s pairwise flux correlations against all other reactions (see **Methods**)^32^. Biologically, this metric has been used in analyses of cancer metabolic models to identify therapeutic targets that destabilize the cancer cell’s metabolic networks and functioning^32^, and we recently demonstrated that high RCC reactions align with central genes in a diatom’s gene co-expression network^30^. *Second*, we compared RCC density distributions between each TOGSMM (**Supplementary Fig. 14**) to compute a community-wide per-sample ‘synergy rank’ score (see **Methods**). This score, derived from the comparison of RCC distributions, investigates a phenotype that cannot be derived from metagenomic comparisons. We interpret this score to represent the biochemical interdependence of the modeled superorganism such that enrichment for low RCC reactions lowers the synergy rank (i.e., closer to zero) with the implication that few reactions control the system, and reaction fluxes are more independent from others as compared to samples of higher synergy rank. This score thus evaluates mechanistic independencies between metabolic reactions under the thermodynamic and redox constraints. It corresponds to an emergent trait that we posit could represent (loosely here due to the current superorganism-assumption requirement) the level of community-wide metabolic reaction synergy and/or cooperativity (**Fig. 2a**). Together these new metrics help describe each seawater sample’s metabolic phenotype, which we in turn used to study as modeled biogeochemistry processes occurring in each sample as follows.

## Metabolic synergy across the ocean

Starting from the community-wide synergy rank metric, we next explored how it related to metadata throughout the global oceans dataset (**Fig. 2b**). This revealed associations between synergy rank and abiotic factors such as iron concentrations (estimated from the NEMO-PISCES Earth System Model, Spearman r: -0.5, p-value: 2×10^-5^), which are known to impact plankton communities^33^ and, now, our findings would suggest also impact per-sample metabolic phenotypes. Alpha diversity of mOTUs (i.e., a biodiversity taxonomic metric) was also significantly associated with synergy rank (Spearman r: 0.27, p-value: 1×10^-3^). While the correlation is weak (e.g., putatively due to the lack of metabolic autotrophic system), we interpret it to inform the “diversity-stability debate”^34^ whereby synergy rank indicates community metabolic network stability, such that in a high synergy system, high RCC reactions are more prone to perturbation-driven instability than in a low synergy system. If future work confirms this, synergy rank could be used by researchers and policymakers to evaluate ecosystem resilience and stability from the new mechanistic perspective of community-wide metabolism.

Per-sample synergy ranks also revealed biogeographical patterns (**Fig. 3a** and **3b**, and **Supplementary Text 3**). For example, the metabolic phenotypes of most oligotrophic waters, such as the Mediterranean Sea and the Indian Ocean, had low synergy ranks. In contrast, those from nutrient-enriched water masses that follow bloom decay (e.g., Stations 76 and 138) or nutrient release via island effects (e.g., Stations 122 to 125) had high synergy ranks. Such patterns could emphasize a relationship between metabolic synergy and microbial productivity via nutrient availability^35^. Indeed, standard GSM approaches^36^ allow to compute a maximum theoretical growth rate associated with each TOGSMM which is the production of biomass evaluated through the flux through the biomass reaction (See **Methods**). This estimation revealed its correlation with metabolic synergy rank (**Fig. 3c**; Spearman r: 0.70, p-value: 10^-14^), emphasizing the effect of a high synergetic system on theoretical productivity.

**Figure 3.**
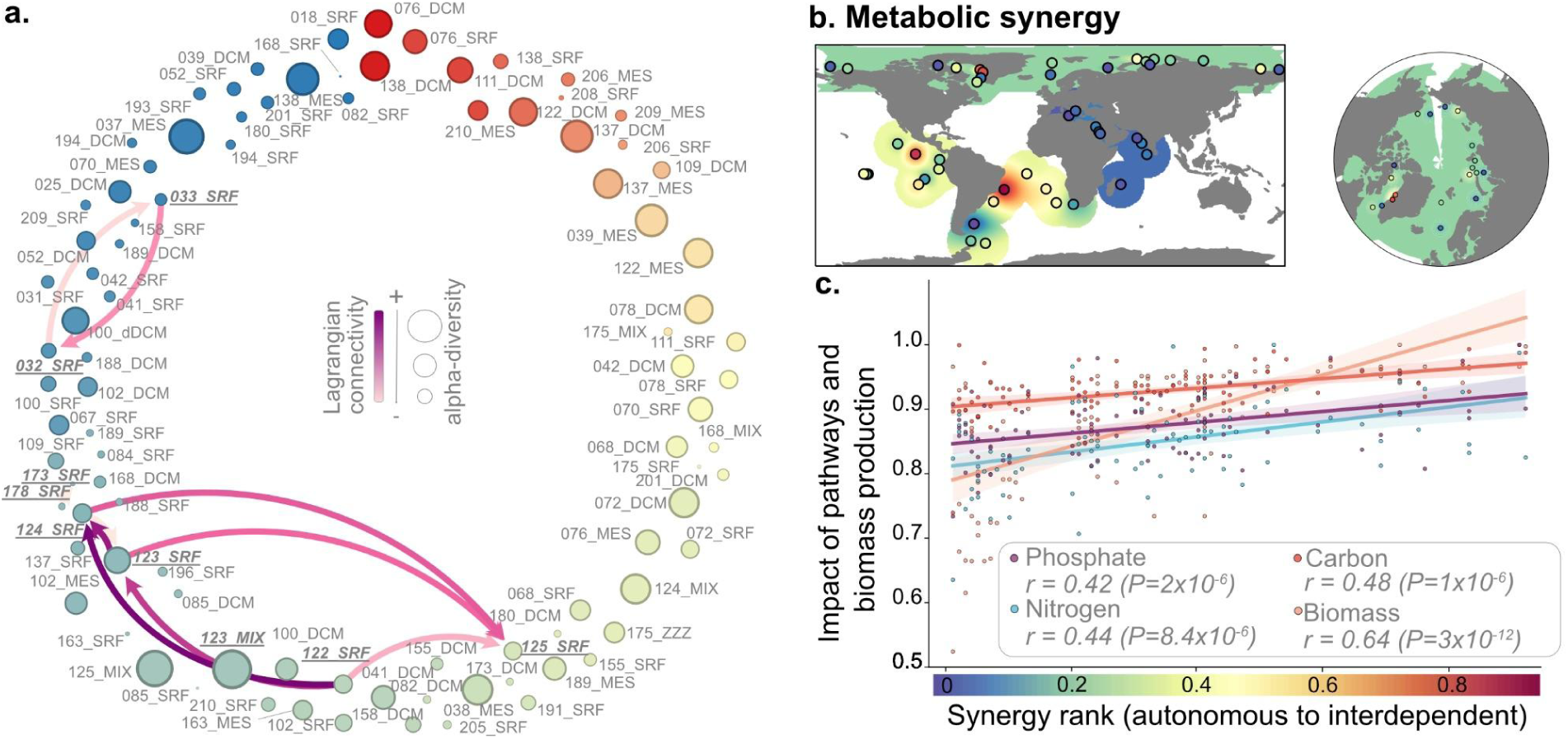
Metabolic synergy across the global ocean. **a.** Distribution of *Tara* samples based on metabolic reaction synergy from autonomous (blue) to interdependent (red) systems. The node size is proportional to the per-sample prokaryotic organism diversity. Directed edges represent Lagrangian connections between samples realized in less than two months to describe the potential acclimation of water masses. The darker the arrow’s color is, the stronger the Lagrangian connection is. In other words, the darker the arrow is, the higher the connection probability between the water masses is. **b.** Projection on the metabolic synergy across the global ocean visualized from an equator- (left) and pole- (right) centric perspectives. The data points were interpolated at each station (halo color on each map), and actual data values are the colored dots. **c.** Relationships between the metabolic synergy of the microbial ecosystem against the normalized impact of phosphate (purple), nitrogen (blue), and carbon (red) reactions and the potential prokaryotic ecosystem biomass production (orange).

Beyond these global observations, we also explored local metabolic features, where higher-resolution sampling was available to test, at the metabolic level, the assumption that most Earth System Models make of a continuous transition between conditions (i.e., each point on the grid)^37^. Specifically, we used Lagrangian connectivity between each TOGSMM (computed as done previously^38^) to ‘connect’ samples to assess how metabolic phenotypes changed over short time scales (60 days or less; see edges in **Fig. 3a** and **Methods**). Though only one sample pairing could be tested (natural fertilization and island effect near the Marquesas islands^33^), it revealed an uneven shift across the synergy rank gradient rather than seeing a continuous transition from an more autonomous state to a more synergetic state (or vice versa) as a water mass is transported (**Fig. 3a** and **Supplementary Fig. 6**). As more high-resolution sample sets become available, it will be ideal to assess their metabolic phenotypes and synergetic status across representative water masses and environmental gradients to better guide ESM assumptions.

Considering another context, we asked whether summed RCCs for biogeochemically relevant reactions (pathways in the KEGG database^39^ for carbon, nitrogen, and phosphorus) offered functional patterns of interest. Such estimations would indicate the relative mechanistic importance of each biogeochemical cycle to sustain the water mass metabolic phenotype. While the summed RCC of each cycle scales with synergy rank (**Fig. 3c** and **Supplementary Text 4.c** for discussion on the statistical validity of this correlation), we also observed a remarkably constant proportional summed RCC (std < 0.1 of the mean value for the cycles C:N, C:P and P:N). These observations require further explorations and improved underlying reaction-to-pathway assignment data (**Supplementary Text 4.a**). Still, they could mechanistically contribute to the debate about elemental ratios in the ocean’s plankton and seawater^40^.

Finally, we evaluated the use of expressed transcripts (grouped into 8,803 KO^8^ categories) against the synergy rank across large ecological gradients (synergy ranks that shifted from positive to negative between polar and temperate subsystems) to assess possible transcript-based biomarkers for the synergy rank metabolic phenotype. Indeed, some KOs showed significant correlations (**Supplementary Fig. 13**) with expected functions involved in core metabolism (i.e., oxidoreductase activity) or membrane activity (i.e., transferase, molecular transducer, membrane transport, and transmembrane transporter activity). However, surprisingly, this also revealed functions including hydrolases and lyases, which are commonly encoded by viruses. We interpreted this latter signal to suggest that viruses impact community-level metabolic phenotypes and sought to assess this signal further by shifting to more virus-targeted assessments.

## Virus-modulated metabolisms and shunt/shuttle biogeography

To explore the role of viruses in modulating community-level metabolic phenotypes, we looked at the presence of conservatively-identified, virus-encoded AMGs^42^ using viral metagenomes within TOGSMMs. Strikingly, this revealed that virus-encoded AMGs do not randomly target reactions, but instead predominantly target the highest RCC reactions needed to sustain the community metabolic phenotype (**Fig. 4a**, and **Supplementary Fig. 14**). Biologically, viruses are known to obtain host genes at random through error-prone genomic replication and individual-level selection that genomically fixes those that improve viral fitness^43^. However, these findings show that this evolutionarily selective process also manifests at a community-level such that viruses target high RCC reactions. These biochemical reactions demonstrate topological centrality within metabolic networks and enable fluxes that regulate flux distribution across the entire range of reactions. Consequently, they are pivotal to the overarching metabolic phenotype of the community and serve as a foundation for its stability, predicting that a perturbation on their flux would drastically impact the whole.

**Figure 4.**
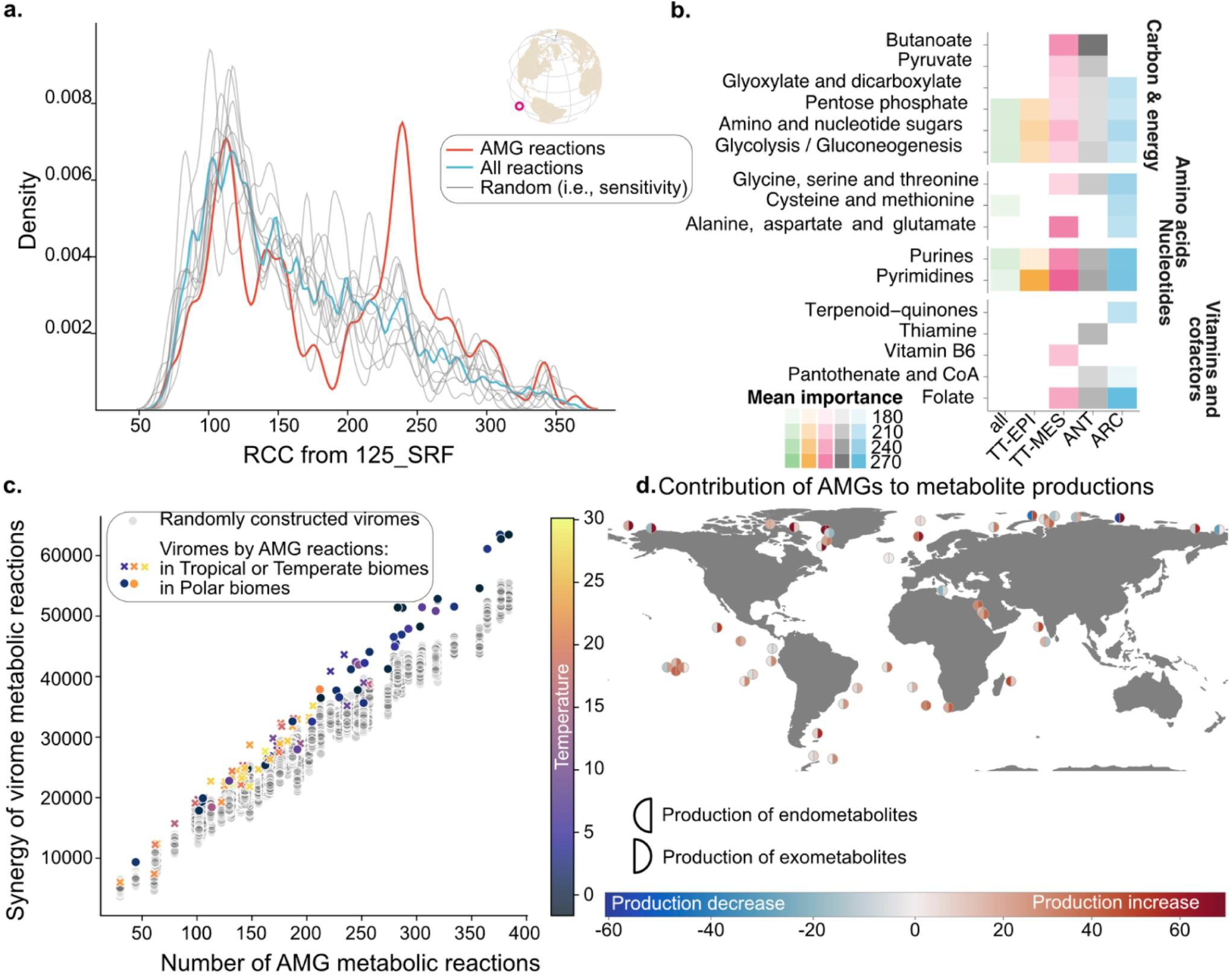
RCC and impact of reactions targeted by virus-encoded Auxiliary Metabolic Genes (AMGs). **a.** Representative example from sample 125_SRF of the RCC of all (blue) and randomly sampled (gray, see **Materials and Methods**) metabolic reactions as compared to the reactions targeted by virus-encoded AMGs (red). All samples showed a similar, statistically significant high RCC AMG spike (see **Supplementary Fig. 14)**. **b.** Metabolic pathways associated with high RCC AMG-targeted reactions across ecological zones. The RCC of reactions targeted by AMGs, present in all networks and statistically belonging to the high RCC spike, were averaged per KEGG pathway (i.e., lines) and ecological zone (i.e., columns). RCCs are then displayed by opacity. Note that a single reaction may be shared among several KEGG pathways. **c.** Comparison between the summed RCC of reactions targeted by AMGs and the number of reactions per sample. Each dot represents a sample. Its color code indicates the temperature associated, and its shape characterizes the system: a circle for polar systems and a cross for temperate systems. Grey dots represent the summed RCC of randomly defined viromes. **d.** Contribution of AMGs to exometabolites and endometabolites through increased (i.e., red) or decreased (i.e., blue) metabolite production. Here we consider metabolites that are only exometabolites (n=26), or only endometabolites (n=28)^41^.

Exploring this further functionally, we found that virus-encoded purine and pyrimidine AMGs – already known to be enriched in viral metagenomes^44,45^ and differentially expressed in marine virocell experiments^46^ – were conserved in all TOGSMMs, but were among the highest RCC reactions in TOGSMMs specific to different ecological zones (**Fig. 4b**). Presumably, this reflects their critical role in the energy demand of viral replication^47^, but now TOGSMM-contextualization suggests nucleotide-synthesis AMGs also significantly impact the community-level metabolic phenotypes. Other energy-related AMGs targeted fermentation, glycolysis, gluconeogenesis, the pentose phosphate pathway, and two-carbon compound reactions. Presumably, such AMG-impacted metabolisms arise from varied viral host cell reprogramming responses across gradients of nutrient-impacted host physiological states requiring diverse energy resources to drive the viral life cycle. Again, while these kinds of functions are characteristic from virocell experiments^42,43^ and beginning to be explored under *in situ* conditions^48,49^, TOGSMM-contextualization now places them into a mechanistic metabolic modeling framework that intertwines cellular systems-level functionality with ocean basin and even global ecological context. For example, AMG-targeted amino acid metabolism genes varied with low-energy cost amino acids (glycine, serine, threonine, alanine, aspartate and glutamate)^50^ targeted broadly, higher energy cost amino acids (cysteine and methionine)^50^ were exclusively targeted in the Arctic Ocean (**Fig. 4b**). Given that cysteine and methionine are involved in sulfur metabolism and sulfur-related AMGs are distributed globally in the ocean^51^, their polar enrichment suggests polar-specific impact of viruses on the salvage of amino acids for nutrient acquisition. Further, AMG reactions were significantly more abundant and associated with higher RCC in polar systems (**Fig. 4c**), which suggests a stronger impact of viruses on polar community metabolic phenotypes^52^. Together, such TOGSMM-powered findings highlight critical metabolic pathways for viral replication that can help constrain virocell studies and metabolic models of viral infection while also placing them into a broader biogeochemical and biogeographical context.

Finally, we wondered whether we could use TOGSMMs to predict virus ecosystem “shunt versus shuttle” effects, which is of collective interest ^41^ due to the ocean’s climate buffering role. For a quarter-century, the paradigm was that viruses help heterotrophs rapidly remineralize carbon by making it more available via cell lysis – i.e., the viral shunt hypothesis^53^. However, aggregating lysis products^54,55^ and viral roles in carbon cycling beyond lysis (e.g., AMGs and metabolic reprogramming) have emerged as additional contributors to biogeochemical cycling^56^. At least in some instances, viruses are now thought to ‘shuttle’ carbon to the deep sea via sinking aggregated cells and/or altering grazing efficiency^48,53,57,58^. Accordingly, previous *Tara* Oceans work using statistical taxon-based modeling suggested that virus abundances, more than those of prokaryotes or eukaryotes, best predict key biogeochemical processes like carbon flux^52^.

Expanding upon these prior findings, we used TOGSMMs to explore per-sample shunt-versus-shuttle emphasis interpreted through the lens of the relative contribution of AMGs previously categorized as exo- or endo-metabolites^41^. For this analysis, we assumed that (i) exometabolites represent sticky materials that increase carbon export via aggregation invoked in the shuttle hypothesis^59^ and (ii) endometabolites, as intracellular metabolites, are instead linked to community-level reprogramming and, thus, the remineralization-related shunt hypothesis. These are strong and highly over-simplified assumptions due to poorly characterized seawater metabolites^41^ and the need to more broadly characterize metabolic reactions into shunt-versus-shuttle effects by better assessing biological products’ stickiness^59^ at scale. Visualizing epipelagic layer AMG reactions revealed estimates per-sample shunt-vs-shuttle proportions and possible biogeographic hot-spots (**Fig. 4d**, and **Supplementary Fig. 15** for details), where this currently imperfect estimator (due to the myriad input data issues) already revealed an overall larger ‘shuttle’ contribution consistent with the oceans currently being a carbon sink via particle aggregation^59^. Even given these assumptions, our mechanistically modeled exometabolite predictions weakly but significantly correlate to Particulate Organic Carbon (Spearman r: 0.34, p-val 4×10^-3^). This suggests promise for global evaluation once better input data arise in the form of an ideal ‘metabolite stickiness’ list, a resource the genome-scale modeling community could use immediately to improve predictions.

## Conclusions

This study introduces a mechanistic modeling approach to abstract community-wide heterotrophic bacterial metabolism from ‘omics data, and uses a statistical approach to explore the resultant per-sample metabolic phenotypes. Just as genome-scale models have revealed at a cellular level how biological systems self-organize to reveal metabolic linchpins and emergent cellular properties^60^, our approach reveals how, from a given seawater sample’s metabolic capacity, one can establish ‘metabolic phenotype’ throughout the global oceans, and metabolic reaction essentiality for each sample’s community metabolism. This found that biotic and abiotic factors can influence variation in global ocean metabolic phenotypes, while also providing new hypotheses about viruses underpinning community-wide metabolic stability and a framework for assessing their roles in the planet-critical ocean biological carbon pump. Future developments will have to incorporate autotrophic reactions, ‘omics abundances, and/or genomic content refined from MAGs to broaden the set of research questions that can be addressed, but this new mechanistic modeling framework already offers an omics-powered approach to help transform our understanding of how microbiomes and viromes drive ecosystem functioning.

## Materials and Methods

### The global ocean dataset

The dataset for metabolic network reconstruction consists of paired prokaryote- (n = 127) and virus-enriched (n = 127) metagenomes and the corresponding prokaryotic metatranscriptomes (n = 94) from previously published datasets from *Tara* Oceans (TO) and *Tara* Oceans Polar Circle (TOPC) expeditions^8,26^. These samples were collected from 62 sampling stations, spanning the global ocean, including the South Pacific Ocean, North Pacific Ocean, South Atlantic Ocean, Indian Ocean, Red Sea, Mediterranean Sea, Southern Ocean, North Atlantic, and Arctic Ocean.

### Generation of the reference gene catalog

Representative coding sequence (cds) or gene features were selected for prokaryotic microbial and viral fractions. In the case of prokaryotic cds features, the Ocean Microbial Reference Gene Catalog (OM-RGC.v2)^8^, which included 46,775,174 features, was used as the reference catalog. In the case of viral cds features, genes were called from assemblies in virus-enriched metagenomes (n = 127) using Prodigal (v2.6.1) in the metagenomic mode^61^. The resulting genes were grouped into clusters using mmseqs2^62^ (easy-cluster, -s 7.5, e-value 0.001) at two sets of parameters (30% ANI across 70% of the length and 60% ANI across 80% of the length). To establish functional relevance, 1000 gene clusters, with more than 20 genes per cluster, were randomly selected for each set of parameters. For functional annotation, the cds features within each cluster were searched against the RefSeq database (v208). Ambiguous clusters were further assessed using the Pfam database^63^ (v33.1). Collectively, this analysis identified representative cds features (n = 70,347,973) clustered at 30% ANI across 70% of the length as the reference catalog for subsequent modeling efforts.

### Calculation of metagenomic and metatranscriptomic profiles

We profiled metagenomic and metatranscriptomic cds features for use in constructing TOGSMMs. The gene and transcript abundance calculations of OM-RGC.v2 from the prokaryotic metagenomes (n = 127) and metatranscriptomes (n=94) were previously reported^8^ and used in this study. In total, 26% of the 47M cds features in OM-RGC.v2 were mappable to metabolic reactions in the BiGG database^64^ (v1.6) and were thus considered in TOGSMMs. To calculate gene abundance from viral metagenomes, amino acid sequences of representative cds features were first searched against the BiGG database^64^ (v1.6) using DIAMOND^65^ (v0.9.24.125) (--more-sensitive, e-value 0.001). This resulted in 7% of the 70M cds features (n = 4,939,042) matched to reactions in the BiGG database^64^ (v1.6). Quality-controlled reads of viral metagenomes^26^ (n = 127) were then non-deterministically recruited to the 5M cds features using CoverM (v.0.2.0) (https://github.com/wwood/CoverM) with the trimmed mean method. Reads were kept if they were with ≥ 95% identity and ≥75% read coverage. Genes with < 70% coverage within a sample were not considered. Similar strategies were used to calculate the metatranscriptomic composition and map prokaryotic metatranscriptomes (n = 94) to the cds features in the viral metagenomes.

### Building a *Tara* Ocean Genome-Scale Metabolic Model

Metabolic networks were generated per paired samples. For each paired sample, genes with an abundance > 0 calculated from metagenomes were piped through CarveME^25^ (v1.5.1) using a universal model of bacterial metabolism. The resulting metabolic networks (n = 127) were gap-filled using the marine media (**Table S1**). Similar strategies were applied to genes with transcriptomic abundance > 0. These identified 94 networks derived from the transcriptomic abundance profile.

We used the classical constraints used in metabolic modeling to obtain the model. To each network, we associated a set of constraints:

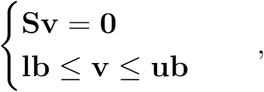

where S ∈ ℝ*^m,n^* is the stoichiometric matrix of the network composed of *m* metabolites and *n* reactions, v ∈ ℝ*^n^* is the flux vector that represents the flux that goes into each reaction of the network, and lb, ub ∈ ℝ*^n^* are the lower and upper bounds of the flux vector (default values from the CarveMe algorithm). TOGSMMs are available here: https://uncloud.univ-nantes.fr/index.php/s/9nXG6kRz66CwDAW (a zenodo link and a DOI will be added upon acceptance of the manuscript).

Each TOGSMM is prone to Flux Balance Analysis. Under the superorganism assumption, the estimated solution of the heterotrophic prokaryotic growth objective function (the biomass reaction) is interpreted as the theoretical maximum growth rate at the ecosystem level. While not quantitatively accurate, estimating these theoretical maximum growth rates is sufficient to compare ecosystems built following the same procedure.

### Auxiliary metabolic genes and their mapping in TOGSMMs

Systematically cataloged AMGs from the global ocean were used to assess the role of viruses in community-level metabolic phenotype. The Global Oceans AMG catalog was previously established. Briefly, AMGs were called on viral contigs with additional curation steps to exclude non-viral regions, mobile genetic elements, and other genes that may function in the random integration of microbial metabolic genes^42^. To map the Global Oceans AMG catalog (n = 22,779) in TOGSMMs, we searched them against the BiGG database (1.6) present in the CarveMe universe using DIAMOND (v2.0.14.152, –more-sensitive). We then applied the logical rules in CarveMe’s GPR to obtain the reactions AMG supports.

### Sampling of the *Tara* Ocean Genome-Scale Models

Each metabolic model defines a solution space *F*, that is, the ensemble of solutions that satisfies the constraint:

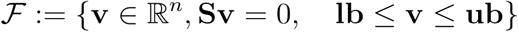

Describing the solution space of a metabolic model is equivalent to assessing its metabolic phenotype. To do so, we used optGpSampler^31^, which allows us to draw random flux distributions from the solution space. We constrain the biomass reaction to produce at least 10% of its maximum. This subjective threshold allows us to discard states with low biomass production, focusing our attention on states that assure the ecosystem’s survival.

We sampled 10^5^ points with a thinning of 10^4^. From this sampling procedure, we computed a pairwise Pearson correlation between each network reaction, resulting in the correlation matrix C ∈ *R*^(*n,n*)^.

### Reaction Cumulative Correlation and cycles

Reaction Cumulative Correlation (RCC) is the sum of its pairwise Pearson correlation in absolute value over the entire network. Formally, we write the reaction’s *j* RCC as

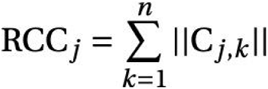

where the sum is done over the set of reactions. A reaction with high RCC is significantly correlated with many other reactions in the network; thus, a change in its flux value will likely change the fluxes of many other reactions. On the other hand, a reaction with low RCC can see its flux vary without changing the overall flux distribution. In graph theory, when abstracting each reaction as a node and each correlation as a weighted edge between nodes, the RCC is called the “strength” of the node. It is equivalent to the centrality of its corresponding multigraph^66^.

Cycle summed RCC is the sum of the reactions’ RCC that compose the cycle. We used KEGG identifiers available in the CarveMe reaction database to obtain the composition of cycles. To avoid subjective thresholds on correlation values, we used all correlations. The low correlations due to random effects appear in all the reactions and are uniformly distributed. As such, they do not impact our results.

### Viruses impact the system through AMGs

AMGs encoded by viruses have been demonstrated to modify host metabolism sufficiently to alter host biogeochemical/metabolite flux profile^67^. To incorporate viruses in our metabolic network, we considered AMG-targeted reactions in the established TOGSMMs. We estimated the impact of viruses through AMGs using the previously computed reactions’ summed RCC. To compute the impact of viruses on the production of metabolites, we computed the contribution of each reaction in the production or consumption of the considered metabolites. We then examined the impact of an increase in each reaction flux from the reactions supported by AMGs. This impact was computed by multiplying the participation of the reactions to the dynamic of the metabolites by their correlations to the reactions supported by AMGs.

Formally, for a given TOGSMM, this gives:

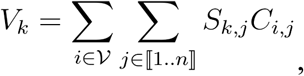

where *ν* is the set of metabolic reactions supported by AMGs, *S_k,j_* is the stoichiometric coefficient of metabolite *k* in reaction *j*, and *C_i,j_* is the pairwise correlation between reaction *i* and reaction *j*.

To estimate random viromes for a given sample, we took the total number of reactions supported by the computed virome and randomly chose that many reactions in the network. This resulted in a random distribution of RCC and a random summed RCC score of reactions targeted by AMGs.

### Stability of the ecosystem

In our framework, stability represents how sensitive the system is regarding a change of flux values. The correlation value from the sampling work allows us to assess this stability. A network with many reactions’ correlations near zero is considered to represent a stable ecosystem because a change in a single reaction’s flux will not significantly change the overall flux distribution and, thus, the estimated behavior of the ecosystem. However, a network with many highly correlated reactions will be more sensitive to changes in a given reaction’s flux, and thus will represent a less stable ecosystem.

We used a Kolmogorov-Smirnov test (KS-test) on the absolute pairwise correlation values distribution to compute this metric. We calculated the KS distance and its associated p-value. We then created the directed graph, where edges were considered if the test p-value was not significant (>5*1e-2), meaning that we could not reject the null hypothesis. An edge from station A to station B means that the ecosystem captured in station A is more stable than the one modeled in station B.

From this graph, one can compute each node’s out-degree, which is the number of edges that start with the considered node. Once normalized, this out-degree corresponds to the sample’s synergy score, as represented in **Fig. 2**.

### Endo and exo-metabolite identification and linking to reactions

Endo and exo-metabolites were determined using Table 1 of Moran et al 2022^41^. The metabolites table was joined to the reagents and products of each reaction used in the models to determine which reactions were directly associated with the transformation of endo and exometabolites.

## Acknowledgments

Our survey was made possible by the sampling and sequencing efforts of the *Tara* Oceans Project. We are indebted to all who contributed to these efforts and other open-source bioinformatics tools for their commitment to transparency and openness. *Tara* Oceans (which includes the *Tara* Oceans and *Tara* Oceans Polar Circle expeditions) would not exist without the leadership of the *Tara* Oceans Foundation and the continuous support of 23 institutes (https://oceans.taraexpeditions.org/).

## Author contributions

D.E designed the experiment. E.SG computed the Lagrangian data. F.T reconstructed the models. A.R simulated the models. A.R, F.T and G.S analyzed the data. A.R, F.T, G.S, MB.S and D.E wrote the initial manuscript. J.R, CM.A-G, P.B, M.P, C.HV, A.L, E.SG, C.T, L.G, SJ.H, D.I, E.K, and A.M help writing the final manuscript.

## Competing interests

SJH has a conflict of interest; he is a co-founder of Koonkie Inc., a bioinformatics consulting company that designs and provides scalable algorithmic and data analytics solutions in the cloud.

This article is contribution number XXX of *Tara* Oceans.

## Data and materials availability

All data used in this paper is already available (-omics) and the models used will be released upon acceptance of the manuscript.

## List of Supplementary Materials

- Supplementary Text 1- 7
- Supplementary Fig 1-15 (Supplementary Fig. 14 is in a separate PDF)

