## Supplementary Materials for "Planetary-scale heterotrophic microbial community modeling assesses metabolic synergy and viral impacts"


#### Supplementary Text 1: Bioinformatics pipeline

A

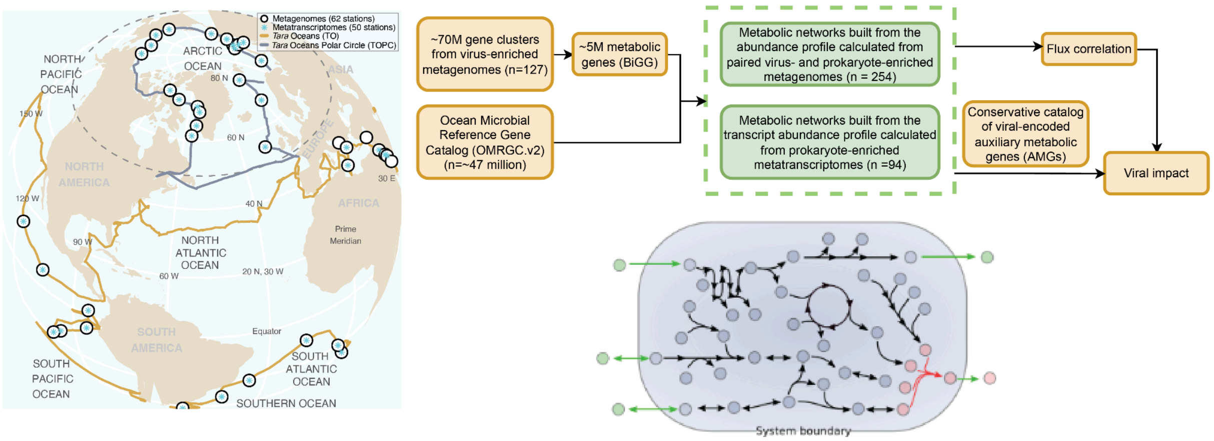


**Supplementary Fig. 1 Pipeline for the metabolic network reconstruction of TOGSMMs**

### Supplementary Text 2: Analysis between Metatranscriptomic- (metaT) and Metagenomic-based (metaG) *Tara* Ocean Genome-Scale Metabolic Models

#### a) Inclusion of the metatranscriptome into the metagenome

The hypothesis of the inclusion of the metatranscriptome into the metagenome was not enforced during the reconstruction of the metabolic model. We used the same universal model whether we were considering genes or transcripts. However, we observed that this hypothesis is still relatively accurate. Considering the networks as a set of reactions, one can compute the cardinal of the interaction of two paired networks (metagenomic-based and metatranscriptomic-based) and their union. On average, the intersection ratio over the number of reactions in the metatranscriptomic-based (metaT) network is 0.99. However, the mean ratio with the number of reactions in the metagenomic-based network (metaG network) is 0.96. Only one metaT network has more reactions than its corresponding metaG: 122_DCM, with nine more reactions.

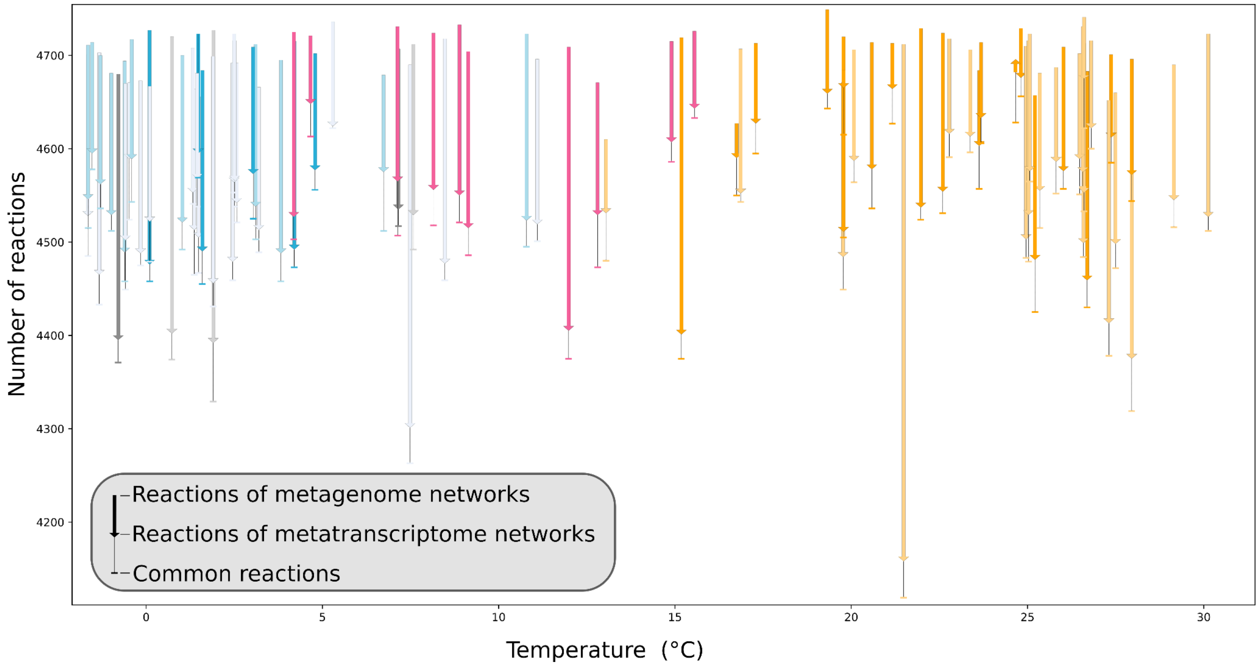


**Supplementary Fig. 2. Description of the paired networks through their reactions.** MetaG networks have a mean of 4,701 reactions (std: 24), while metaT networks have a mean of 4,541 reactions (std: 81)

The difference in the distribution of reactions in metaT and metaG networks could be interpreted as the fact that the water masses have a slow evolving set of metabolic genes (in terms of presence/absence). However, its transcriptome is more variable, reflecting different adaptations of the prokaryotic community to various environmental conditions.

#### b) Rationale for studying metaT networks

As previously mentioned, metaT-based networks are more variable and, as such, can represent different systems across the global ocean. To confirm this hypothesis, we analyzed their "core" metabolic reactions (reactions present in every network) and compared them by calculating Jacquard distances. We confirmed our hypothesis by showing that the core set of metabolic reactions is also smaller in metaT networks than in metaG, even though there are more metaG networks (see **Supplementary Fig. 3**).

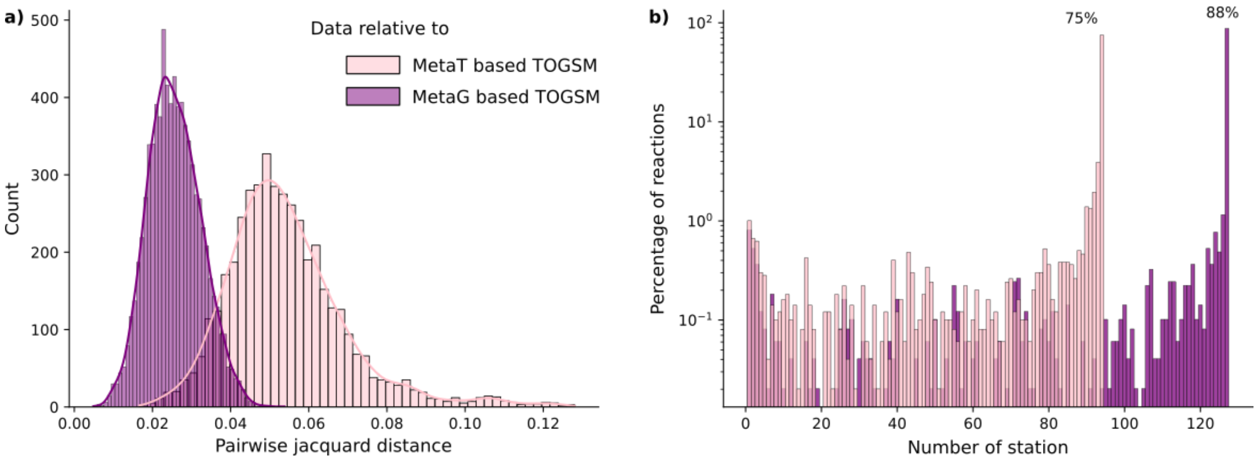


**Supplementary Fig. 3. Topological differences between metaT and metaG based TOGSMM. a)** Distribution of pairwise distance (Jacquard) when networks are considered as sets of reactions. **b)** Distribution of the reactions among the stations. 75% of the reactions are present in all the networks based on metaT, and 88% are in all those based on metaG.

The set of metabolic reactions present inmetaG and metaT networks allow for the recovery of previously defined ecological zones[1] (see **Supplementary Fig. 4**). However, ecological zones are less structured when the importance of the reactions are considered. This indicates that the importance of reactions incorporates a notion of survival in the data. Indeed, while sampling the metabolic phenotype of the water mass, we included a constraint on the biomass reaction to force it to produce at least 10% of its maximum. We thus removed all the potential solutions with less biomass production. It would be interesting to explore how this evolves if we sample with a stronger or weaker constraint on the biomass.

[1] Gregory, A. C. *et al.* Marine DNA Viral Macro- and Microdiversity from Pole to Pole. *Cell* **177**, 1109-1123.e14 (2019).

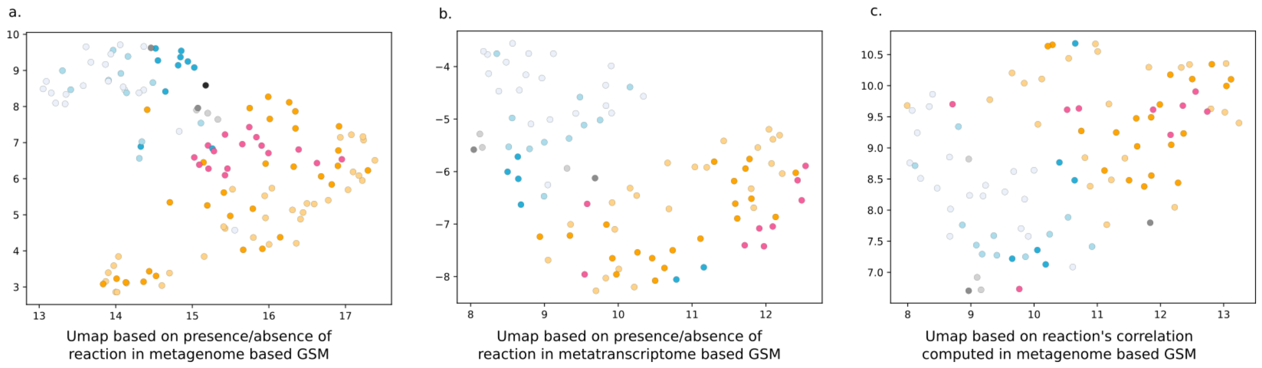


**Supplementary Fig. 4. Uniform Manifold Approximation and Projection (UMAP) of different types of data. a)** UMAP on the presence/absence of metabolic reactions in metaG networks. **b)** UMAP on the presence/absence of metabolic reactions in metaT networks. **c)** UMAP on the importance of reactions in the metaT networks.

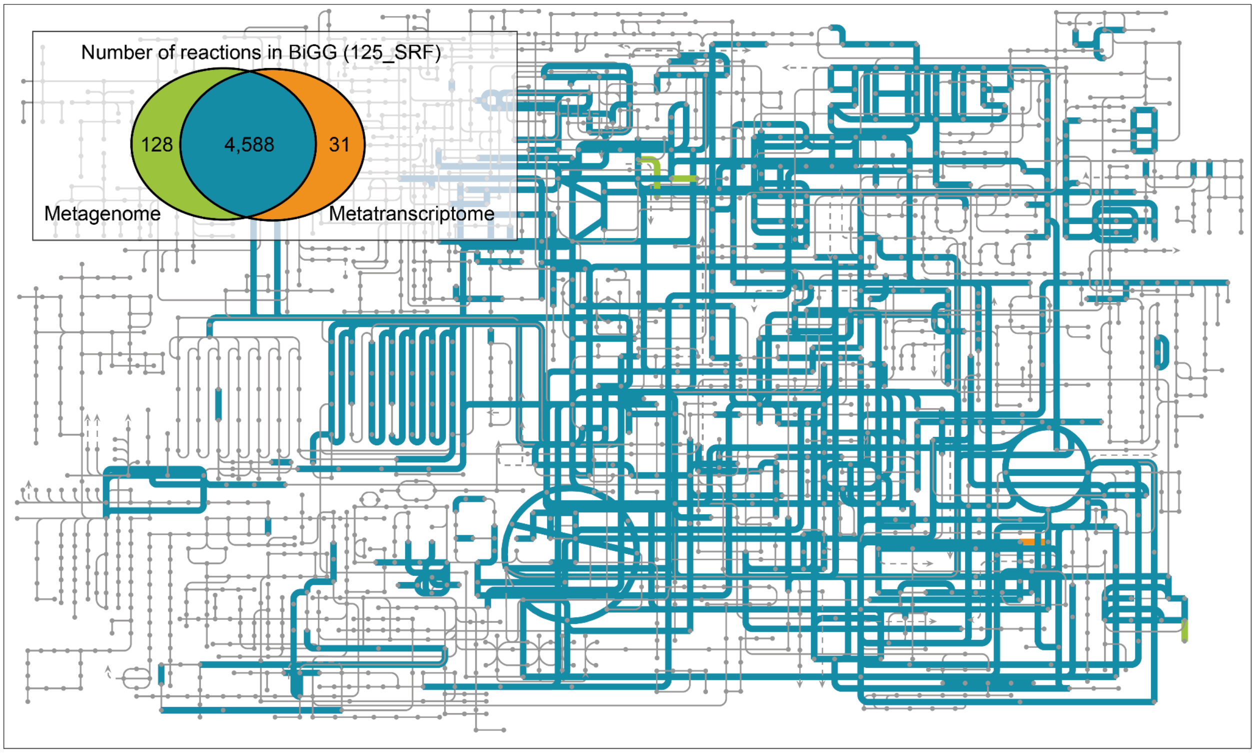


**Supplementary Fig 5**. Metabolic reactions in TOGSMMs derived from the metagenome (green) and metatranscriptome (orange) of 125_SRF.

Supplementary Text 3: Synergy across the oceans and an illustration of mesoscale investigation in the Marquesas archipelago

a) Synergy across the oceans

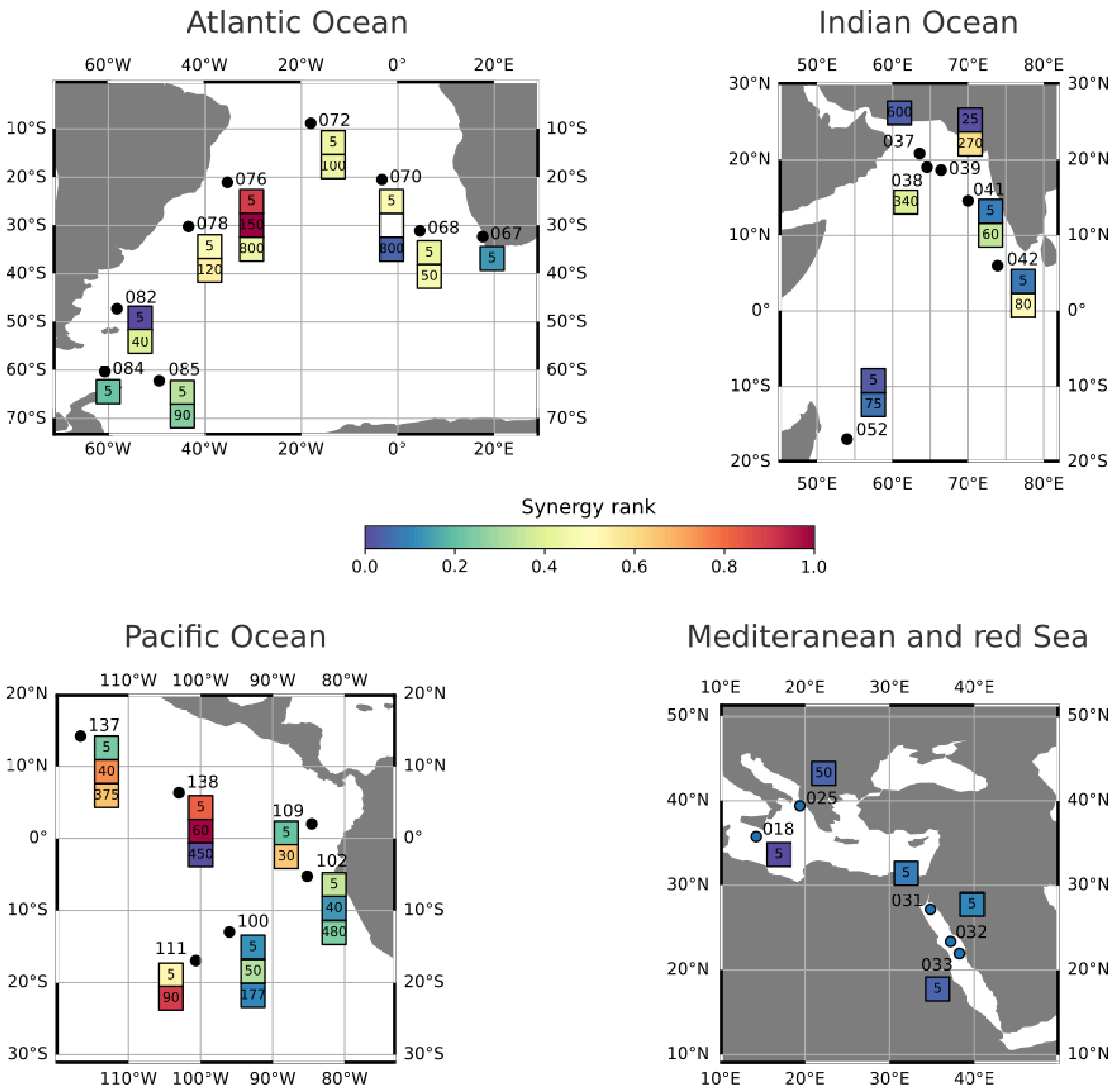

**Supplementary Fig 6. Distribution of synergy rank across different oceans.** For each station, the samples are represented by squares, colored by their synergy rank, and annotated by their depth.

b) Synergy across the Marquesas Islands

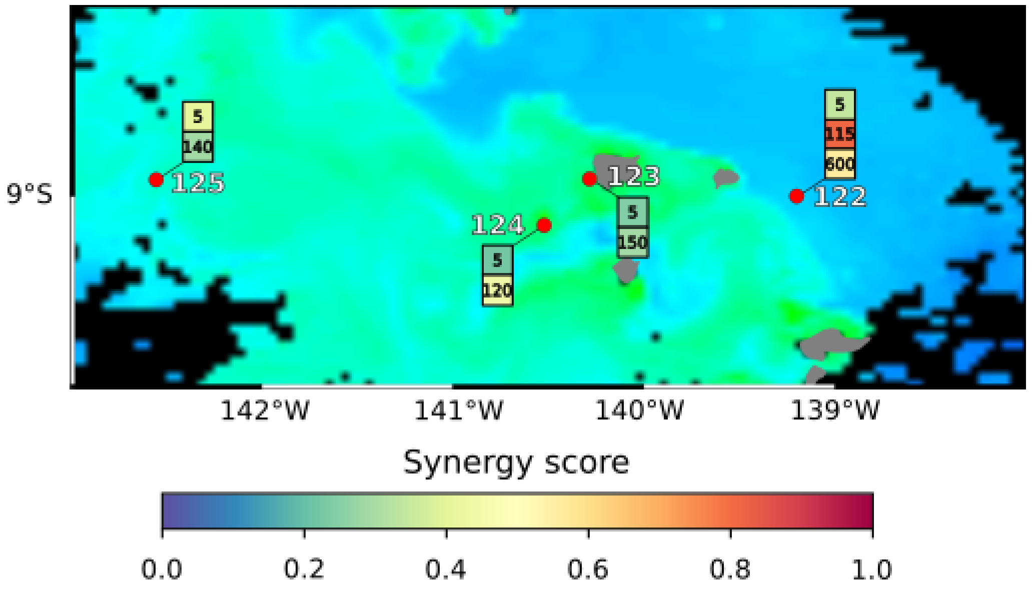


**Supplementary Fig 7. Synergy rank across the archipelago.** The color of the ocean is proportional to the amount of chlorophyll as determined by Aqua-MODIS on the 03/08. Station 122 SRF was sampled on 26/07 and station 125 MIX on 09/08 in the year 2011.

### Supplementary Text 4: a new perspective on Redfield's ratios

#### a) Cycles description and importance

In our dataset, we used the KEGG database to define three cycles: carbon (Carbon metabolism) composed of 100 reactions, nitrogen (Nitrogen metabolism) consisting of 16 reactions and phosphorus (Pentose phosphate pathway) composed of 18 reactions. We also added a phosphorus cycle based on the OrthoPhosphate pathway consisting of 664 reactions (OP cycle). To analyze these cycles, we considered their random definitions: for a given cycle, we sampled $n$ reactions, with $n$ being the cardinal of the considered cycle.

As shown in **Supplementary Fig. 8**, the importance distributions across all networks are consistently above the random cycle importance distributions for KEGG-defined cycles. This effect is less striking for the nitrogen cycle. However, the OP cycle shows the opposite.

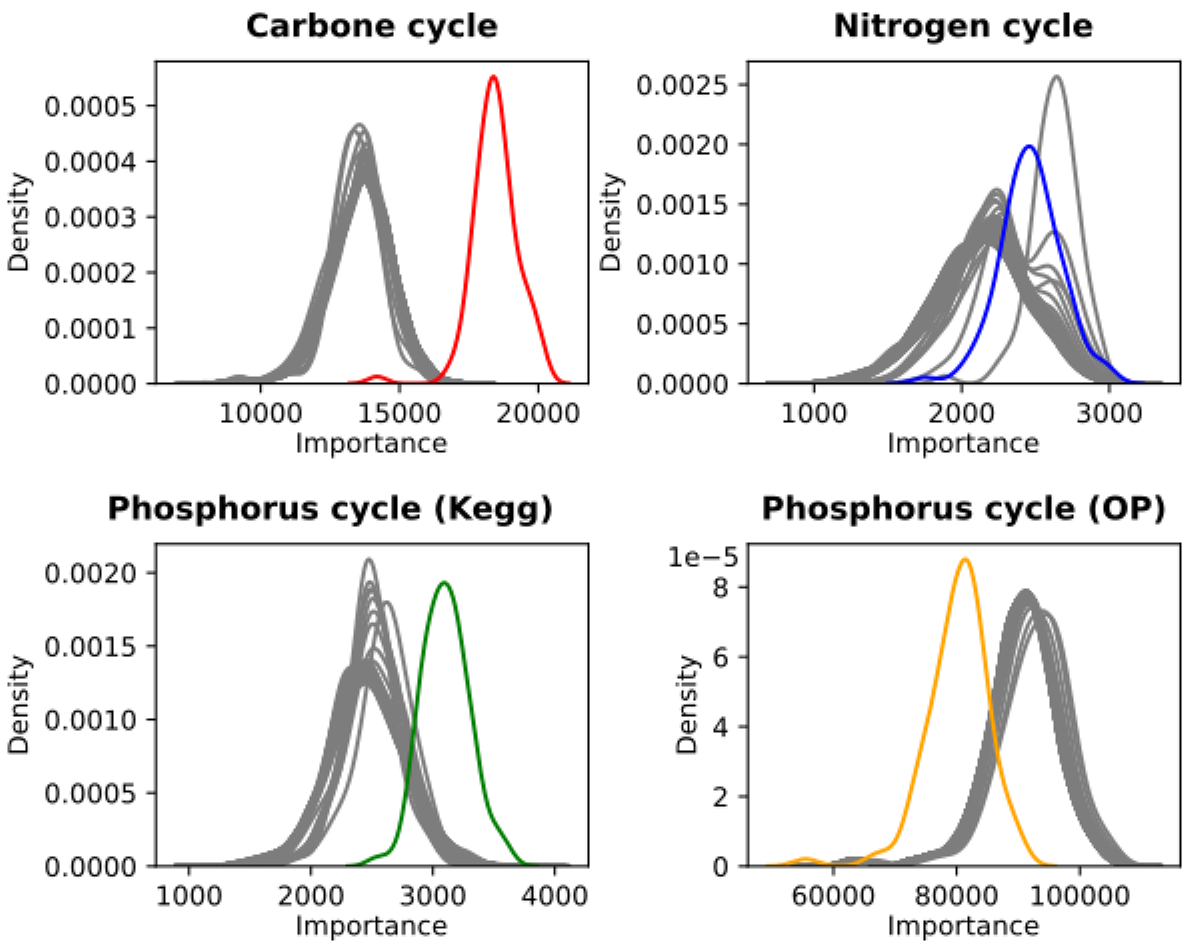


**Supplementary Fig. 8. Distribution of cycle importance.** For each cycle, grey distributions represent randomly chosen cycles (n = 100). Carbone and nitrogen cycles are based on KEGG. Phosphorus is either based on KEGG or contains all the reactions that use the metabolite OrthoPhosphate (OP).

#### b) Redfield ratio and importance ratio

Here, we propose to study reaction importance ratios to reflect the seminal Redfield ratio from a metabolic point of view. We can see the statistical validity of this ratio (see **Supplementary Fig. 9**), as the ratio computed from the data is always on the tail of the two random distributions (one distribution for the numerator, another for the denominator). OP cycles are even “outside” the two distributions (see **Supplementary Fig. 9**).

This analysis shows that the reaction importance ratios are not randomly distributed. When the ratio includes the carbon cycle, we can see that random carbon ratio distributions are far from the actual distribution. The effect is even more substantial when considering the carbon cycle with the phosphorus OP. This effect reflects the non-random distribution of the ratio and the fact that the importance distributions of those two cycles are far from the random distributions. This effect must be clarified when considering nitrogen and phosphorus KEGG cycles due to their relatively small size.

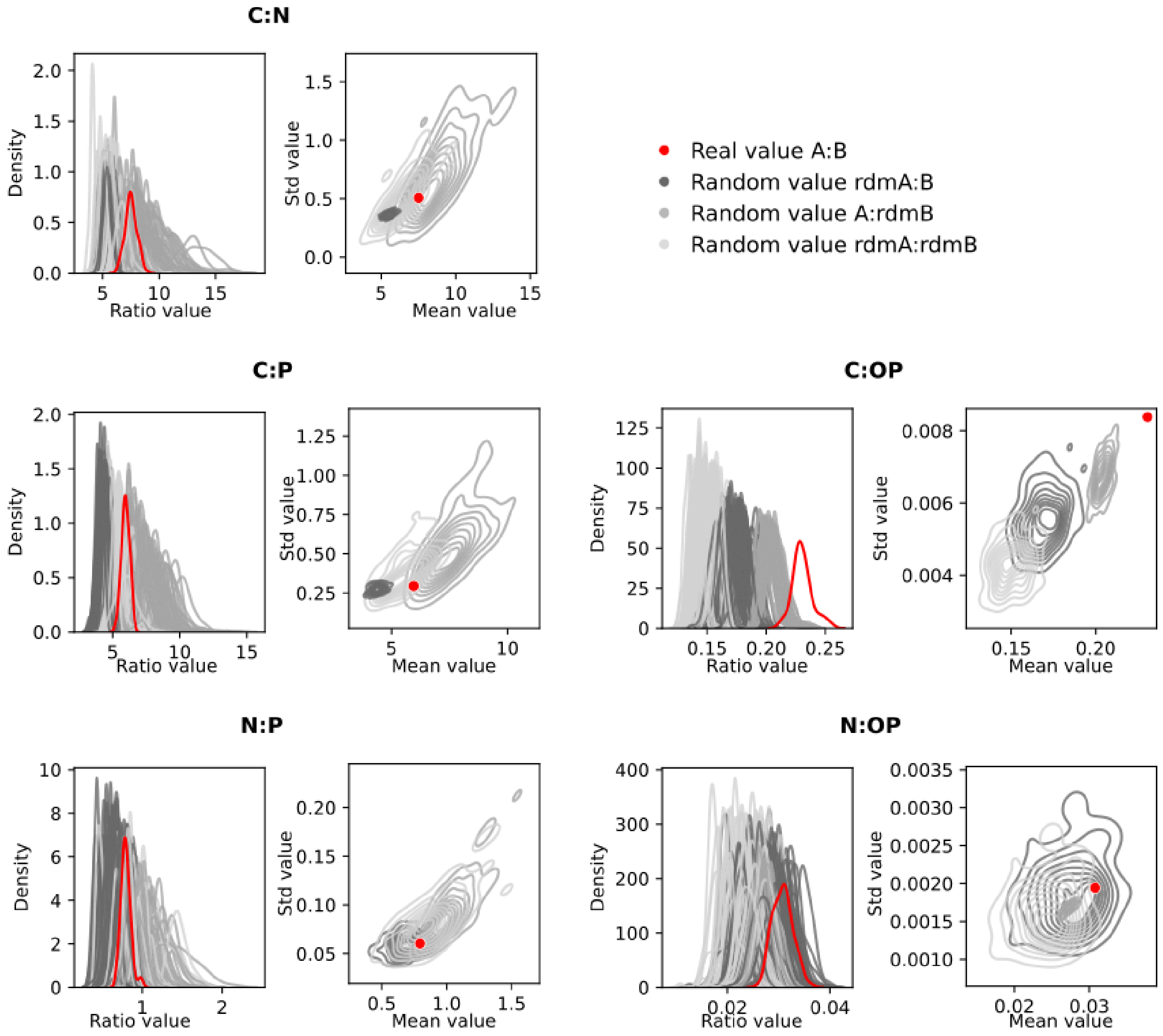


**Supplementary Fig. 9. Distribution of cycle ratios.** The left panel represents the different random ratio distributions (n = 100) for each ratio. On the right panel is the distribution of mean and standard deviation values.

#### c) Correlation of the cycles

We next sought to verify the statistical validity of the correlation of the cycles compared to random cycles. We can see (**Supplementary Fig. 10**) that while carbon and phosphorus OP are statistically different, nitrogen and phosphorus KEGG are almost right at the peak of the distribution. This effect is due to our networks' low representation of the nitrogen and phosphorus cycle. However, we can observe that all correlation values are on the left side of the distributions. It means that cycle importance tends to be more independent of the synergy rank than random cycles. These results also point toward a certain stability of the cycles across all samples.

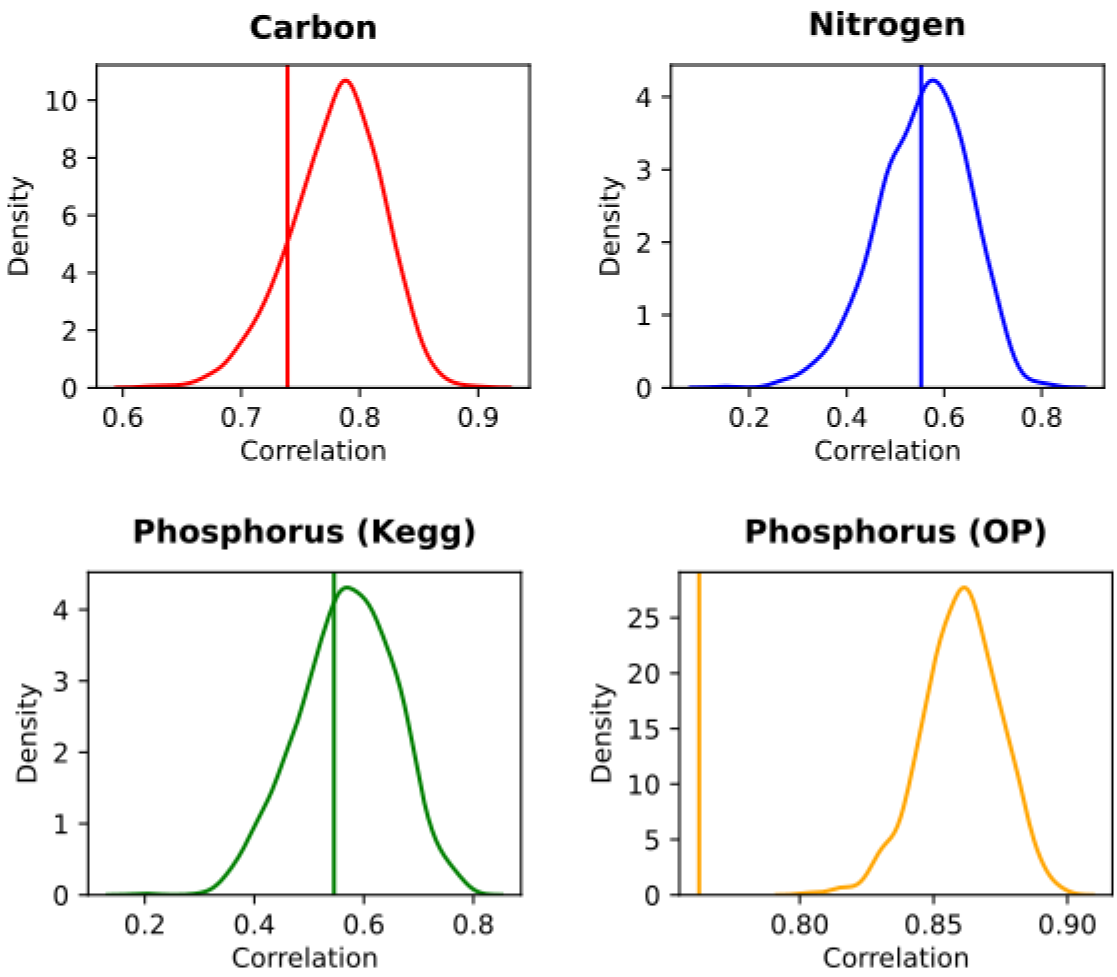


**Supplementary Fig. 10. Distribution of correlation with the synergy rank versus the real correlation.** Vertical lines represent the real Spearman correlation, while the distributions represent the distribution of correlation when using random cycles (n = 1000).

### Supplementary Text 5: Polar and non-polar biomes

#### a) Polar and non-polar biomes distinction

From different types of -omics data, we confirmed the distinction between polar and non-polar biomes. From the whole set of metatranscriptomic to the subset of KO expression, two distinct clusters always emerge, fitting polar and non-polar provinces. This led us to consider how the synergy rank calculated from the polar and non-polar  biomes varied with other parameters.

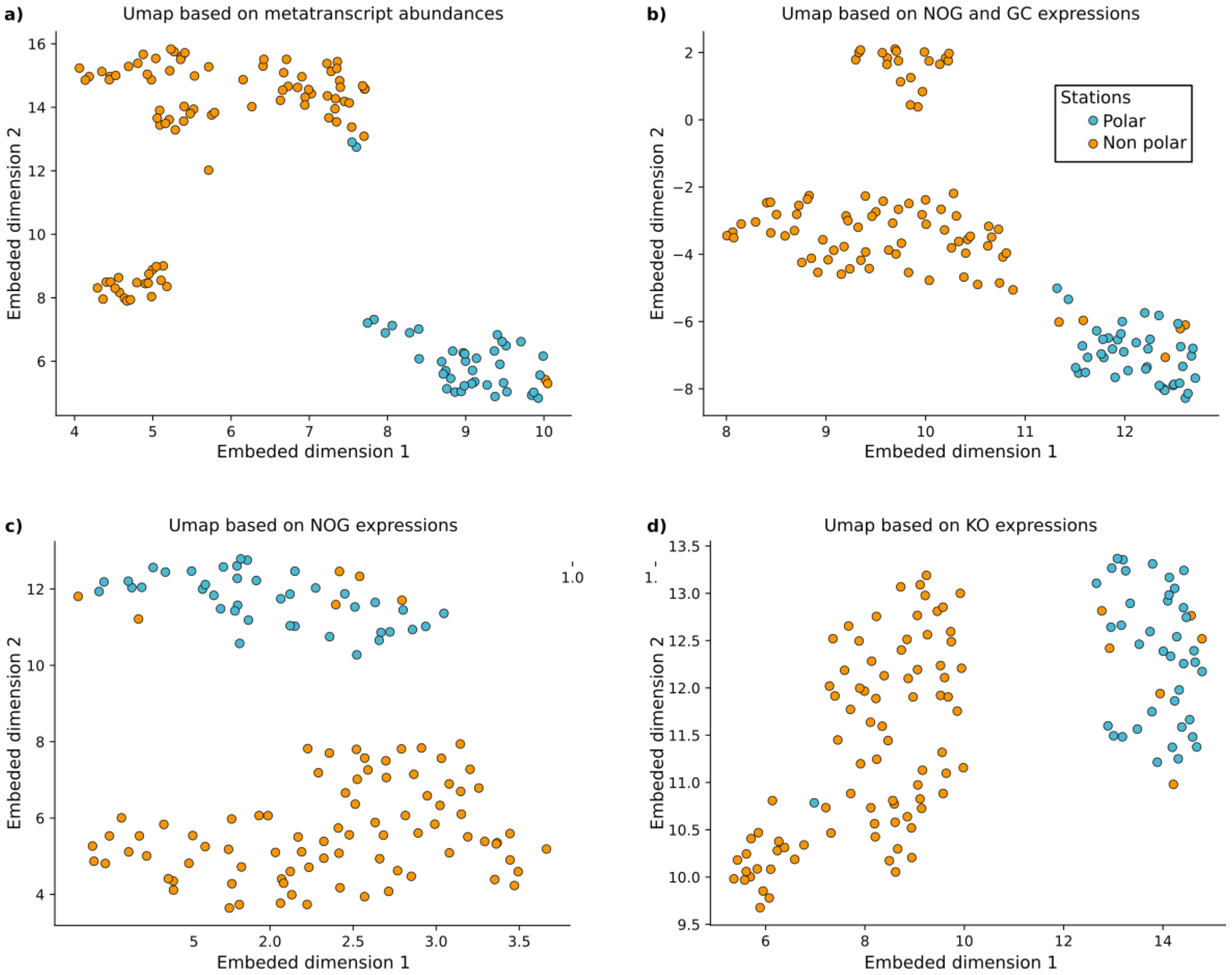


**Supplementary Fig. 11. UMAP on different types of -omic data.** All the different data come from *Salazar et al. 2019*: from the wider emsemble to the most narrow: meta-transcriptomic abundance data, genes not annotated (GC) and genes annotated through eggNOG database (NOG) and genes anntotated through KEGG Orthology (KO).

#### b) Synergy rank vs. different meta-omics data

Given the biogeography of the global oceans identified taxonomically and metabolically (**Supplementary Fig. 11**), we next assessed how metabolic synergy rank related to *in situ* biological data. More correlations are found in polar than tropical-temperate subsystems (**Supplementary Fig. 12**). Focusing on the specific plankton taxa (i.e., meta-barcode) reveals that most eukaryotes associated with high synergy rank were parasitic (i.e., *Syndiniales* in polar subsystem) or less described (i.e., *Heterodinium*), denoting their benefit of living in a synergetic community. For prokaryotes, heterotroph *SAR406 and* autotrophs cyanobacteria and *Rhodospirallaceae* correlated strongly with synergy rank (**Supplementary Fig. 13a**). The former may provide clues as to SAR406's little-understood biogeographic drivers. To deepen our understanding of these systems, we further assessed expressed transcripts (regrouped in KO) that correlated with synergy rank and focused on those that showed synergy rank correlations shifting from positive to negative between polar and temperate subsystems (**Supplementary Fig. 13b**). This revealed genes associated with metabolism (i.e., oxidoreductase activity), membrane activity (i.e., transferase, molecular transducer, membrane transport, and transmembrane transporter activity), and hydrolase and lyase activities.

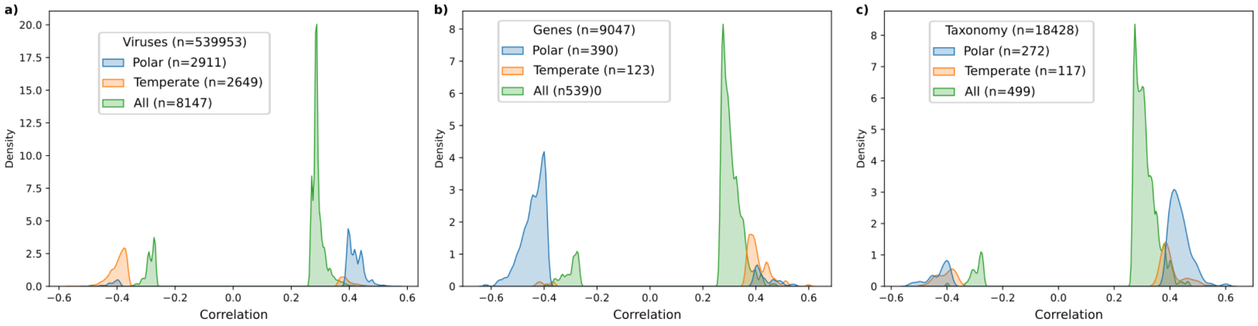

**Supplementary Fig. 12. Correlation of different omic data with synergy rank.** Three systems were considered, polar (blue), temperate or tropical (orange), and the global ocean (green). Correlations are considered if they are with a p-value $p < 0.01$. We correlate the synergy with relative abundance data of viruses (**a**), genes (**b**), and taxonomy (**c**).

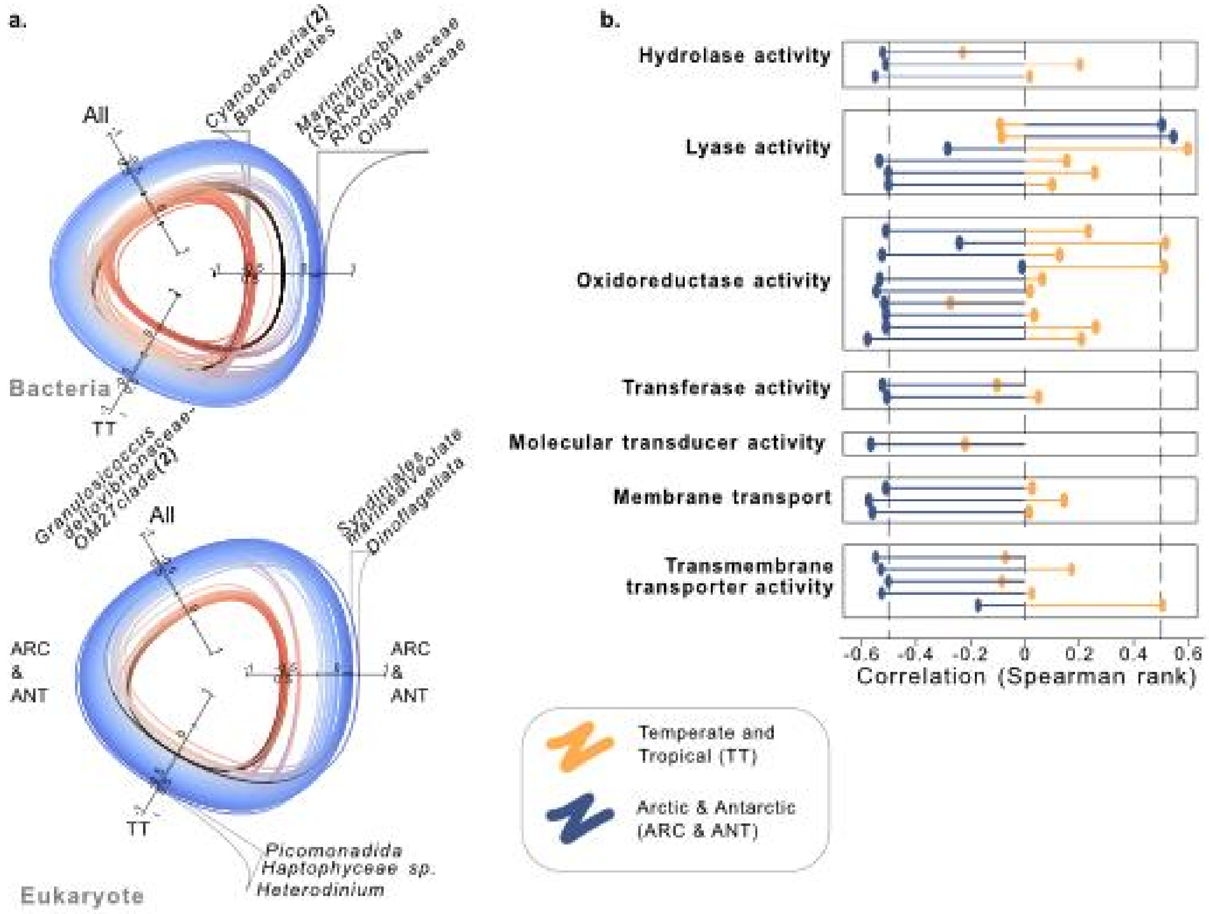


**Supplementary Fig. 13. Identification of major organisms and functions associated with the metabolic synergy rank in temperate or tropical and polar biomes. a.** Hive plots of significant Spearman rank correlations between taxon abundance (separated into Bacteria and Eukaryotes) and synergy rank. Correlations are computed considering different biomes: Polar (i.e., ARC & ANT), Temperate or Tropical (i.e., TT), and global ocean (i.e., All). Each line represents one organism. Blue and red lines indicate positive and negative correlations with the synergy rank, whereas black shows the absence of the considered organism in one biome. Organisms with the highest correlation per biome are emphasized. It stresses the importance of autotrophic and symbiotic taxons. **b.** The highest correlation value of KEGG ID associated with their functions (defined with KEGG and GO annotations) that have significant correlations above 0.5 in one of the arctic and tropical-temperate ocean subsystems.

### Supplementary Text 6: Significative importances in AMGs' reactions

For each TOGSMM, the importance of all reactions fits a Gaussian distribution. This distribution is unimodal when considering all reactions, but bi-modal when looking at the distribution of the importance of the AMGs' reaction. By fitting a mixture of two gaussian models to the distribution of the AMGs' reaction importance, knowing that one of them is the previously fitted gaussian distribution, we can identify the parameter of the second gaussian distribution which corresponds to the AMGs' signal. From the two gaussian models, we then computed the likelihood of an importance value to belong to the first gaussian (the noise) or to the second one (the AMGs' signal). The highest likelihood is used to assign a reaction to the noise or the AMGs' signal.

https://uncloud.univ-nantes.fr/index.php/s/qo5w568mfiCBqP2 (Supplementary Fig.14)

### Supplementary Text 7: Impact of viruses on different metabolites through AMGs

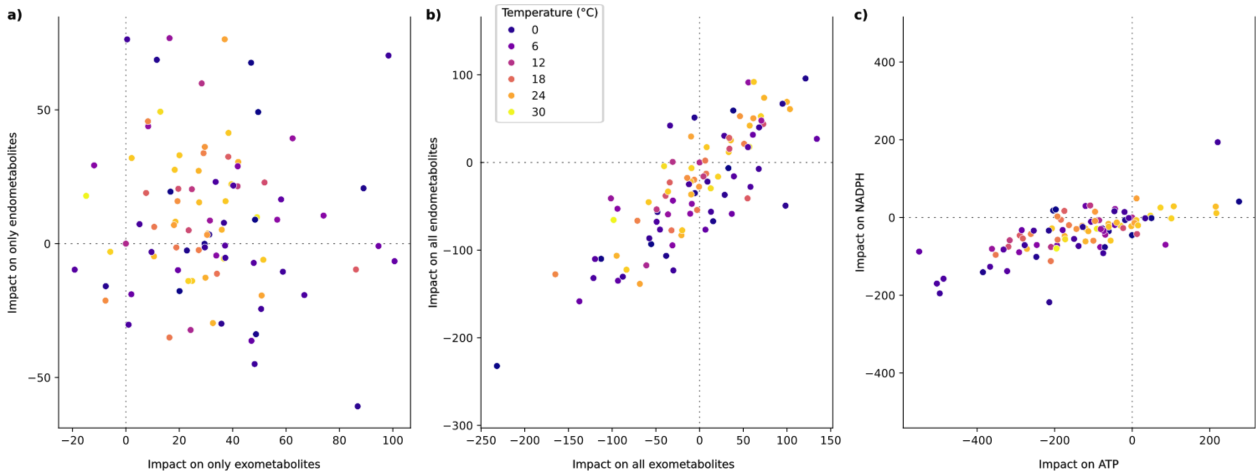


**Supplementary Fig. 15. Impact of viruses on different metabolites through AMGs. a)** Impact of viruses on the production of metabolites that are either exometabolite-exclusive (n = 26) or endometabolite-exclusive (n = 28). **b)** Impact of viruses on the production of metabolites that are either exo- (n = 73) or endo-metabolites (n = 75). 47 metabolites are endo- and exo-metabolites. **c)** Impact of viruses on NADPH production and ATP production.
